# *In Vitro* Activity of Cysteamine Against SARS-CoV-2 Variants

**DOI:** 10.1101/2021.10.02.462862

**Authors:** Jess Thoene, Robert F Gavin, Aaron Towne, Lauren Wattay, Maria Grazia Ferrari, Ranajit Pal

## Abstract

Global COVID-19 pandemic is caused by infection with severe acute respiratory syndrome coronavirus 2 (SARS-CoV-2). Continuous emergence of new variants and their rapid spread are jeopardizing vaccine countermeasures to a significant extent. While currently available vaccines are effective at preventing illness associated with SARS-CoV-2 infection, these have been shown to be less effective at preventing breakthrough infection and transmission from a vaccinated individual to others. Here we demonstrate broad antiviral activity of cysteamine HCl *in vitro* against major emergent infectious variants of SARS-CoV-2 in a highly permissible Vero cell line. Cysteamine HCl inhibited infection of wild type, alpha, beta, gamma, delta, lambda, and omicron variants effectively. Cysteamine is a very well-tolerated US FDA-approved drug used chronically as a topical ophthalmic solution to treat ocular cystinosis in patients who receive it hourly or QID lifelong at concentrations 6 to 10 times higher than that required to completely inhibit SARS CoV-2 in tissue culture. Application of cysteamine as a topical nasal treatment can potentially1) mitigate existing infection 2) prevent infection in exposed individuals, and 3) limit the contagion in vulnerable populations.

## Introduction

Severe acute respiratory syndrome coronavirus (SARS-CoV-2) first emerged in humans during late 2019 in Wuhan, China, and transmitted globally leading to the Coronavirus Disease 19 (COVID-19) pandemic. As of March, 2022, COVID-19 has in resulted in 454,287,804 infections and over 6 million deaths globally. Morbidity and mortality continue to grow due to the emergence of new infectious variants. SARS-CoV-2 is a highly contagious virus transmitted primarily via respiratory droplets. The infection typically results in a wide range of clinical outcome from an asymptomatic state to respiratory failure leading to multiorgan failure and death. While several vaccines have been developed and administered globally, the efficacy of such vaccines against the emergent variants has come into question. Moreover, hesitancy to be vaccinated has also complicated global effort successfully to control the pandemic. Furthermore, antibody-based therapy also has limitations as recently noted for sotroyimab showing decreased efficacy against BA2 subvariant of omicron variant. Clearly, highly effective therapeutic and preventive strategies are needed which, along with vaccines, will control the pandemic.

Here we present data showing *in vitro* inhibition of major currently known SARS-CoV-2 variants by cysteamine. Cysteamine, 2-aminoethanethiol, is a simple aliphatic compound which was first used in man as an antidote to acetaminophen poisoning (1). It was subsequently developed as a treatment for cystinosis after it was found to deplete cultured cystinosis fibroblasts of stored lysosomal cystine, which is the hallmark of cystinosis. Cystinosis is inherited as an autosomal recessive inborn error of lysosomal cystine transport, and characterized chiefly by failure to thrive, progressive renal failure and ESRD by age 10 years (2). In 1994 USFDA NDA approval was granted for cysteamine to treat cystinosis and it also received FDA designation as one of the first Orphan Products (3). Currently, Cysteamine is FDA approved and administered to patients in oral and topical forms to treat systemic and ophthalmic manifestations of cystinosis. Cysteamine is given orally in the systemic treatment of cystinosis. The usual oral dose in children is 50-60 mg/kg/d, and 1.3-1.6g/m^2^/d in adults. These doses yield peak blood cysteamine concentrations of ~ 50-70 μM 60 min after an oral dose (4,5). The corneal crystalline keratopathy of cystinosis is treated with cysteamine eyedrops (6). This treatment is administered 4-8 times per day and continued life-long. Cysteamine has been administered intravenously to two cystinosis patients for 1-10 months (7,8). It is also marketed as a 5% cream used cosmetically to treat melasma (9).

Approximately 800 patients in the United States have nephropathic cystinosis and are on chronic oral and ophthalmic cysteamine therapy, which has become the standard of care since FDA’s 1994 approval. As a genetic disease, cysteamine is required life-long to treat the renal and extra-renal manifestations of cystinosis which include, in addition to progressive renal failure and corneal keratopathy, distal myopathy, neurocognitive disorders, pulmonopathy, endocrinopathy, diabetes, and metabolic bone disease. The general incidence is 1/100,000 live births, although some populations have a higher ratio (10). Prior to approval of cysteamine for treatment of cystinosis, average native kidney survival was <10 years. Currently, with cysteamine, native renal survival can be expected to age 20 years.

Cysteamine has been studied as an antiviral agent. Earlier studies have shown that cysteamine is able to inhibit infectivity of HIV-1 *in vitro* primarily by inhibiting the binding of gp120 with CD4 lymphocytes (11–13). Moreover, recently it has been shown that cysteamine is also capable of inhibiting infectivity of SARS-CoV-2 presumably by inhibiting the binding of S1 protein to ACE-2 receptor (14). Cysteamine and its disulfide cystamine also display an anti-inflammatory effect, reducing SARS-CoV-2 specific interferon-gamma production (15). In this report we examine the inhibitory activity of cysteamine *in vitro* against major variants of SARS-CoV-2 that have emerged so far and find that cysteamine inhibits them at achievable concentrations less than that currently approved by USFDA for ophthalmic use.

## Methods

### Cells and Virus

Vero-TMPRSS2 cells used in the infection assay were provided by the Vaccine Research Center (NIH). The use of this cell line in SARS-CoV-2 infection assay is described elsewhere (16). Cells were maintained in Dulbecco’s modified Eagle’s Medium (DMEM) supplemented with 10% fetal bovine serum (FBS), L-glutamine, 1% penicillin/streptomycin and puromycin (10 μg/ml). Wild type and the variants stocks of SARS-CoV-2 were expanded from the seed stocks in Calu-3 cells by infecting at varying multiplicity of infection in EMEM medium containing 2% FBS, L-glutamine and penicillin/streptomycin. Virus was isolated by harvesting the cell free culture supernatant three to four days after infection depending upon the concentration of the nucleocapsid (NP) protein present in the supernatant as measured by the antigen capture kit (My BioSource). Expanded stocks were shown to be free from the adventitious agent. Identity of each stock was confirmed by deep sequencing. Viruses used in this study include:

Wild type SARS-CoV-2 (P4) isolate USA-WA1/2020 from BEI resources NR-52281; Alpha variant CoVID-19 (2019-nCoV/USA/CA_CDC_5574/2020) from BEI Resources NR-54011; Beta variant CoVID-19 (2019-nCoV/South Africa/KRISP-K005325/2020) from BEI Resources NR-54974; Gamma variant CoVID-19 (hCoV-19/Japan/TY7-501/2021) TY7-503 p1 (Dr. Takaji Wakita, Nation Institute of Infectious Diseases, Japan); Delta variants hCOV-19/USA/PHC658/2021 (B. 1.617.2) from BEI Resources NR-55612 (subvariant 1); NR-55674 (subvariant 2) and NR-55694 (subvariant 3); Lambda variant hCoV-19/Peru/un-CDC-2-4069945/2021 (Lineage C.37) from BEI Resources NR-55656; Omicron variants hCoV-19/USA/MD-HP20874/2021 (Lineage B.1.1.529) from BEI resources NR-56462 (subvariant 1) and from Emory University (17) (subvariant 2).

Infectivity of expanded stocks was determined by plaque forming assay in Vero-TMPRSS2 cells. Infectivity titer (pfu/ml) of each stock used in this study was 3.7×10^7^ pfu/ml for wild type;1.3×10^6^ pfu/ml for alpha; 4.9×10^7^ pfu/ml for beta; 1.8 x10^7^ pfu/ml for gamma;, 2.2 x10^7^ pfu/ml for delta (subvariant 1), 2.9 x10^7^ pfu/ml for delta (subvariant), 2.0 x10^7^ pfu/ml for delta (subvariant 3); 1.8 x10^7^ pfu/ml for lambda; 2.9 x10^7^ pfu/ml for omicron (subvariant 1) and 4.8 x10^6^ pfu/ml for omicron (subvariant 2).

### Cytotoxicity assay

To evaluate the cytotoxicity of cysteamine HCl (ACIC Pharmaceuticals Inc. Brantford, Canada), Vero-TMPRSS2 cells (25000 cells/well) were plated overnight in a 96 well plate in Dulbecco’s modified Eagle’s medium (DMEM) supplemented with 10% FBS, L-glutamine, 1% penicillin/streptomycin and puromycin (10 μg/ml), and the method used in virus inhibition assay was duplicated, using the same concentrations of cysteamine HCl. The cells were preincubated with Dulbecco’s modified Eagle’s medium (DMEM) supplemented with 2% FBS, L-glutamine, 1% penicillin/streptomycin and puromycin (10 μg/ml) (complete DMEM medium) for 2 hours at 37°C in 5% CO2 in 100 μl medium and then transferred to wells containing Vero-TMPRSS2 cells. After 1 hour of incubation, medium from each well was removed and 100 μl of complete DMEM medium containing either no cysteamine HCl or 20% of the original concentration of cysteamine HCl was added to mimic the condition of the infection assay method. The plates were then cultured for 72 hours at 37°C in 5% CO2. Cytotoxicity was measured by adding 100 μl of Cell titer glow (Promega) reagent and incubated for 15 minutes at room temperature. Luminescence endpoint was read in a plate reader (Biotek Cytation 5). Percent cytotoxicity was calculated based on the luminescence reading of cysteamine HCl-treated wells compared to the medium only treated control wells.

### Thiol Quantitation

Measurement of free sulfhydryl concentration in the infection assay—was performed in Vero-TMPRSS2 cells (25000 cells/well) plated overnight in a 96 well plate in Dulbecco’s Modified Eagle’s Medium (DMEM) supplemented with 10% FBS, glutamine, 1% penicillin/streptomycin and puromycin (10 μg/ml). Thiol concentration at the same intervals as cells infected with virus was determined by Ellman’s reagent (5,5’-dithio-bis-2-nitrobenzoic acid, DTNB, Sigma) by reading the optical density at 412 nm in a plate reader (Biotek Cytation 5) as described elsewhere (https://www.bmglabtech.com/ellmans-assay-for-in-solution-quantification-of-sulfhydryl-groups/), and was calculated based on a standard curve generated with the known concentrations of cysteamine HCl.

### Virus inhibition assay

Antiviral activity of cysteamine HCl was assessed in a biosafety level 3 facility under compliance with BIOQUAL’s health and safety procedures using inhibition of virus infection on plaque formation Vero-TMPRSS2 cells (175,000 cells per well) were added into 24 well plates in DMEM medium containing 10% FBS, L-glutamine, puromycin (10 μg/ml) and penicillin/streptomycin and the plates were cultured overnight at 37°C in 5% CO2. For assays measuring dose dependent inhibition of infection, doses of SARS-CoV-2 (see Figure 3) capable of forming measurable number of plaques in the control wells were preincubated with different concentrations of cysteamine HCl at 37°C for 2 hours in a total volume of 600 μl of complete DMEM medium. Cysteamine HCl/virus mixture was then transferred to each well of a 24 well plate of Vero-TMPRSS2 cells in a total volume of 250 μl and incubated for 1 hour at 37°C in 5% CO2. Each well was then overlaid with 1 ml of culture medium containing 0.5% methylcellulose and incubated for 3 days at 37°C in 5% CO2. The plates were subsequently fixed with methanol at −20°C for 30 minutes and stained with 0.2% crystal violet for 30 minutes at room temperature. Plaques in each well were manually scored. Inhibitory potency measured as absolute IC50 was defined as the concentration of cysteamine HCl that resulted in 50% reduction in the number of plaques compared to the untreated controls. The IC_50_ values were calculated using GraphPad Prism 9 program choosing nonlinear regression in a Dose-Response curve. For evaluating the kinetic of virus inactivation both 5 mM and 10 mM concentrations of cysteamine HCl were pre-incubated with SARS-CoV-2 for 0, 15, 30, 60 and 90 minutes and the mixture was then transferred to Vero-TMPRSS2 cells (175,000 cells per well) previously cultured overnight in a 24 well plate in DMEM medium containing 10% FBS, glutamine, puromycin (10 μg/ml) and penicillin/streptomycin. Remaining steps of the infection assay were as described above.

## Results

### Toxicity of cysteamine HCl in culture

Cytotoxicity of cysteamine HCl in Vero-TMPRSS2 cells was determined to select the concentrations to be used in the dose response virus inhibition assay. Cysteamine HCl was preincubated in medium for 2 hours followed by a one-hour incubation with Vero-TMPRSS2 cells. The mixture of cysteamine HCl and medium then was removed, and the cells cultured in fresh medium for 72 hours, following which the viability of the cells was determined as described in Methods. As shown in Figure 1A, no significant toxicity was noted below 20mM cysteamine HCl while some toxicity was noted with 50 mM concentration. To determine if prolonged incubation of Vero-TMPRSS2 with cysteamine as done for the plaque assay would induce cytotoxicity in the target cells, cysteamine HCl was pre-incubated with medium for 2 hours followed by incubation with Vero-TMPRSS2 cells for one hour. The culture medium was then diluted fivefold as done for the plaque assay and the cytotoxicity was measured after 72 hours. Under these conditions, some cytotoxicity was observed at 50 mM concentrations of cysteamine HCl (Figure 1B).

**Figure 1:**
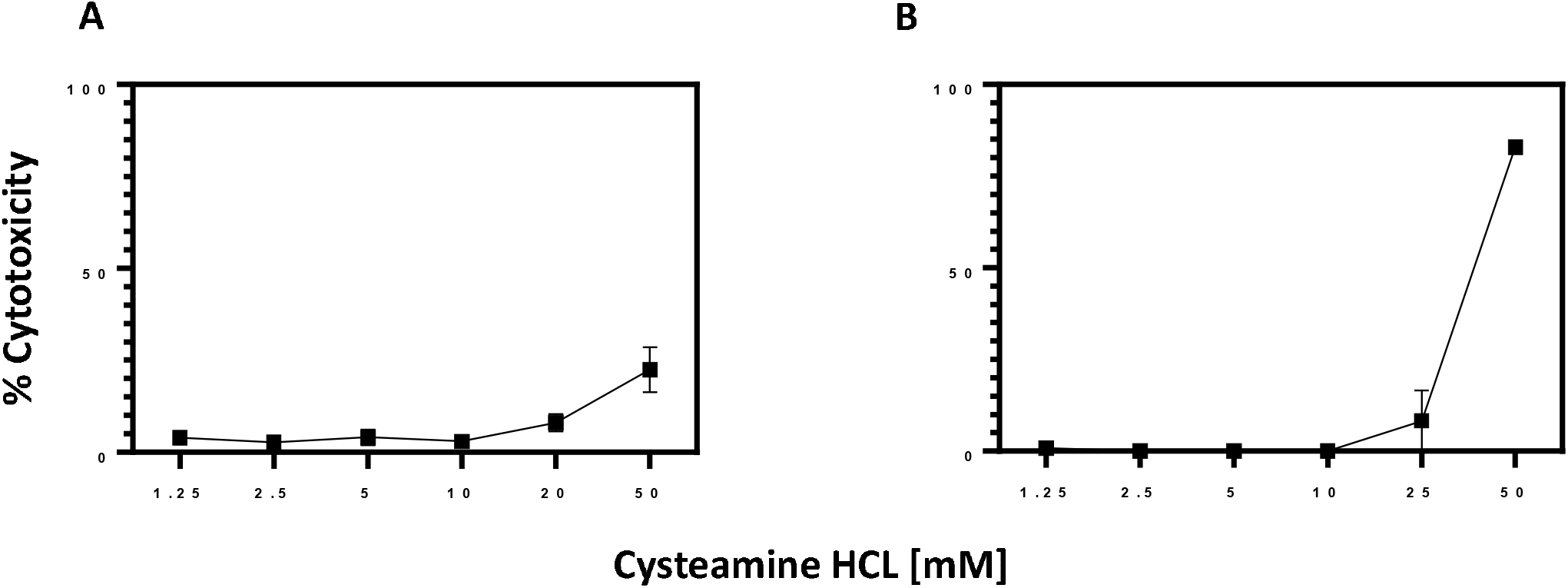
Cytotoxicity of cysteamine HCl in Vero-TMPRSS2 cells. Cysteamine HCl was preincubated with Dulbecco’s modified Eagle’s medium (DMEM) supplemented with 2% FBS, L-glutamine, 1% penicillin/streptomycin and puromycin (10 μg/ml) for 2 hours at 37°C in 5% CO2 in 100 μl medium and then transferred to the wells containing Vero-TMPRSS2 cells. After 1 hour of incubation medium from each well was removed and 100 μl of complete DMEM medium containing either no cysteamine-HCl (A) or 20% of the original concentration of cysteamine-HCl was added to mimic the condition of the infection assay method (B). Cells were cultured for 72 hours at 37°C in 5% CO2 and cytotoxicity measured as described in the Method section.

Because free thiols rapidly oxidize in tissue culture medium, the concentration of free thiol in the culture was measured by Elman’s reagent over a period of 72 h. As shown in Figure 2, there is progressive loss of thiol concentration during this interval. Clearly, no appreciable change in the concentration of free sulfhydryl group was detected both during the pre-incubation of cysteamine HCl with virus for two hours and during the infection step when the virus/cysteamine HCl mixture was incubated with Vero-TMPRSS2 cells for an additional hour. However, a sharp drop in the concentration of -SH group was noted between 24 hours and 72 hours of incubation.

**Figure 2:**
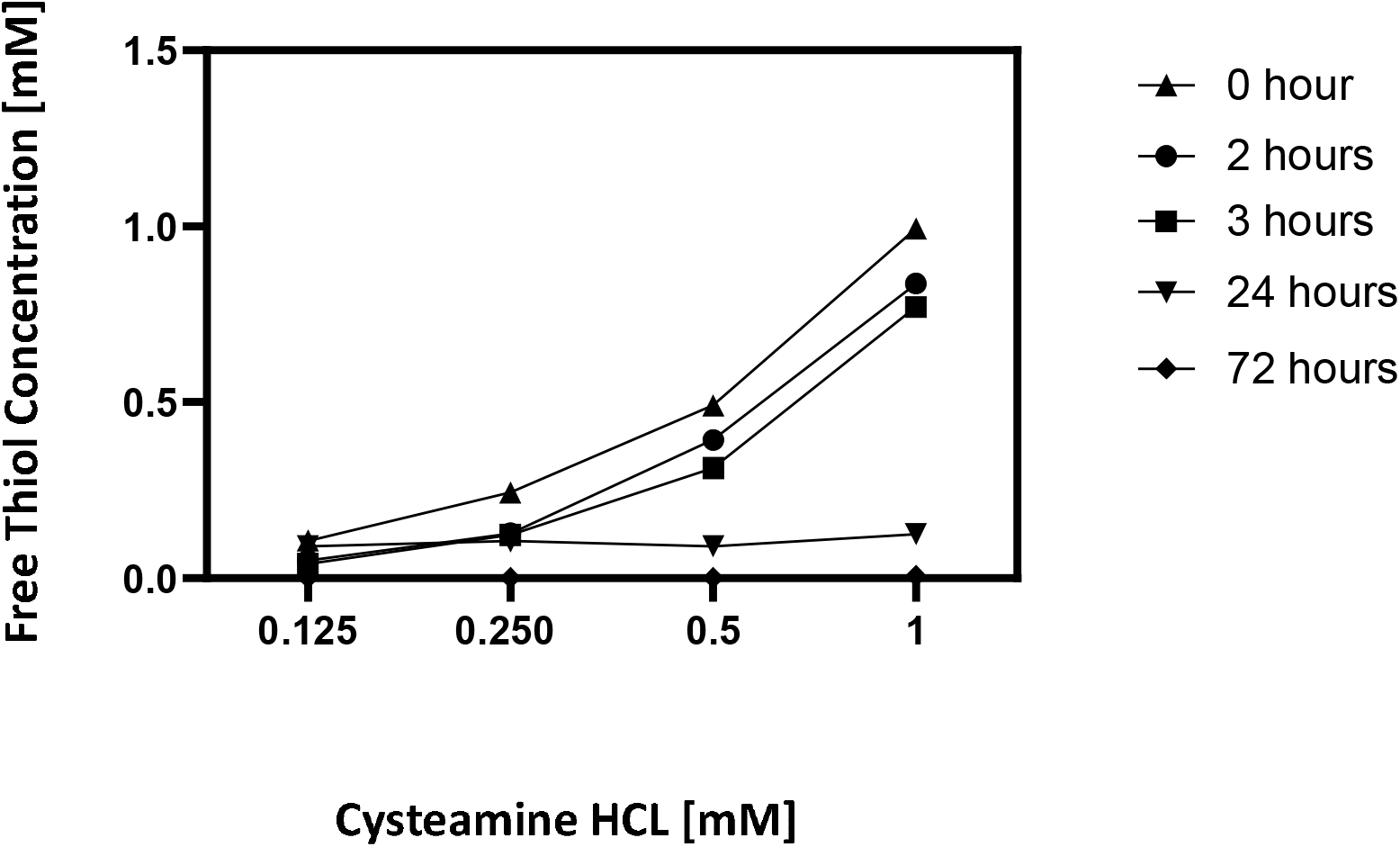
Concentration of free thiol present in the culture: The free thiol content of four concentrations of cysteamine HCl were measured at the initiation of pre-incubation with virus (0 hour), after pre-incubation with virus (2 hour), during incubation of cysteamine HCl/virus mixture with Vero-TMPRSS2 cells 3, 24 and 72 hours post infection as described in the Method Section.

### Inhibition of SARS-CoV-2 variants by cysteamine HCl

To determine dose dependent inhibition of SARS-CoV-2 by cysteamine HCl, Vero-TMPRSS2 was selected, because this cell line is known to be highly permissible to both wild type and the six variants tested in this study. Moreover, these viruses were shown to produce defined plaques when assayed in this cell line (18). As described in the Materials and Method section, SARS-CoV-2 was pre-incubated with different concentrations of cysteamine HCl at 37°C in 5% CO2 for 120 min and the mixture was then added to Vero-TMPRSS2 cells. After one hour of infection the plates were overlaid with 0.5% of methyl cellulose overlay and the plaques were developed and scored after 72 hours. For controls, virus was treated the same way with medium lacking cysteamine HCl. Positive control, using anti-RBD rabbit polyclonal IgG was and assayed in an identical manner. Dose response curves as well as the IC50 values are shown in Figure 3 and Table 1. Wild type WA strain as well as the alpha, beta, gamma and lambda variants were inhibited in a dose dependent manner compared to the control infection with IC50 values ranging from 1.252 mM for wild type to 1.528 mM for alpha (Figure 3A), 0.799 mM for beta, 0.8602 mM for lambda and 1.976 mM for gamma variant (Figure 3B). A similar pattern of inhibition of infection was noted when three different subvariants of delta variant was treated with different concentrations of cysteamine HCl (Figure 3C) with IC50 value of 1.066 mM for subvariant 1, 1.7 mM for subvariant 2 and 1.373 mM for subvariant 3. Similarly, two subvariants of omicron were also inhibited by cysteamine-HCl with IC 50 value of 0.671 mM for subvariant 1 and 1.006 mM for subvariant 2 (Figure 3D). Positive controls using anti-rabbit RBD polyclonal IgG inhibited infection of all viruses in a dose dependent manner (data not shown).

**Figure 3:**
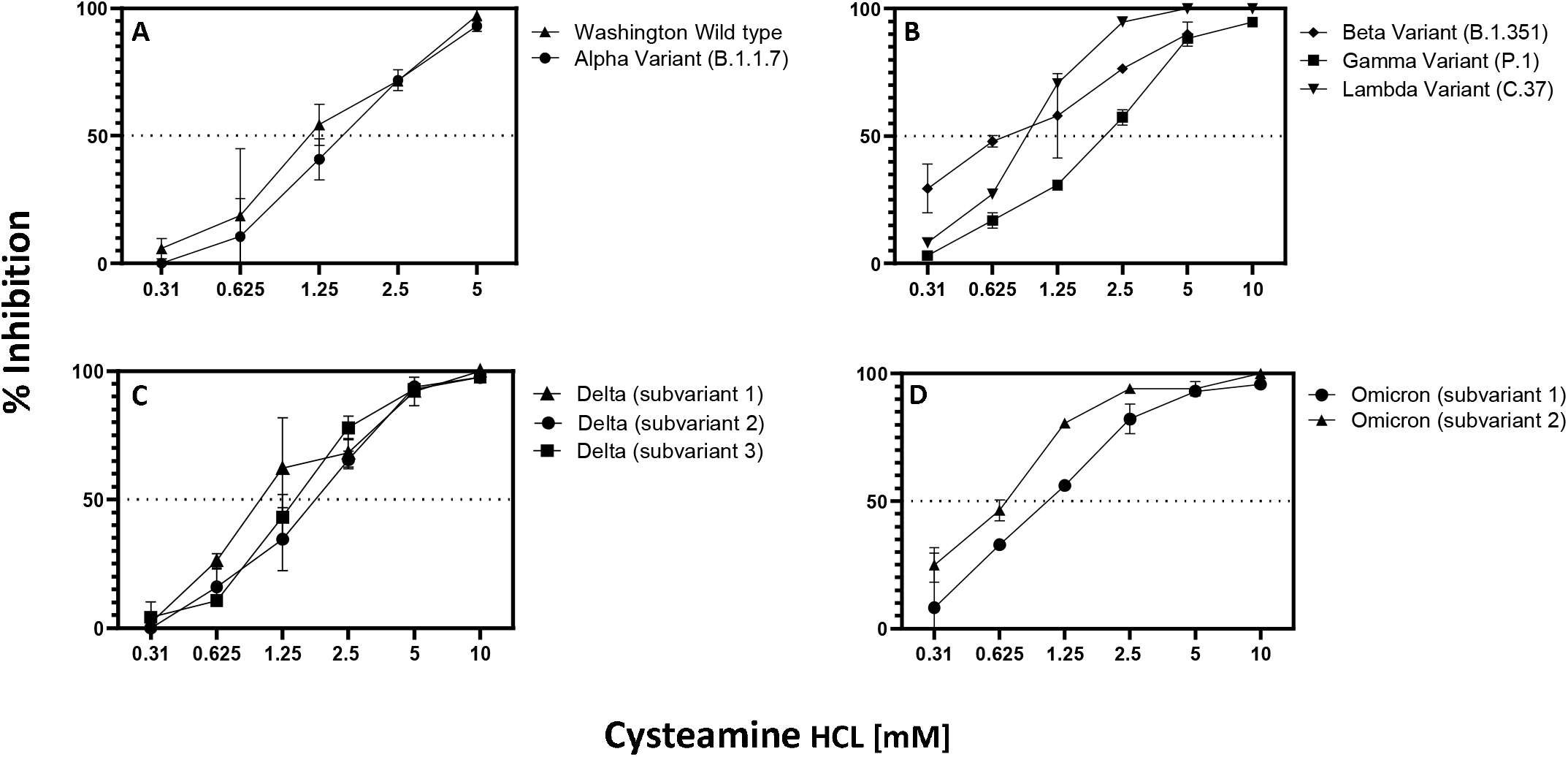
Dose dependent inhibition of infection of wild type and SARS-CoV-2 variants by cysteamine HCl: (A) Wild type (25 pfu/well), alpha (125 pfu/well); (B) beta (18.75 pfu/well), gamma (25 pfu/well); lambda (60 pfu/well); (C) delta subvariant 1 (37.5 pfu/well), subvariant 2 (48pfu/ml), subvariant 3 (42 pfu/ml); (D) omicron B1.1, subvariant 1 (16.8pfu/well) and subvariant 2 (24pfu/well). Virus was preincubated with varying concentrations of cysteamine HCl at 37°C for 2 hours in a total volume of 600 μl of complete DMEM medium as shown in the figure. Cysteamine HCl/virus mixture was then transferred to each well of Vero-TMPRSS2 cells in a total volume of 250 μl and incubated for 1 hour at 37°C in 5% CO2. Each well was then overlaid with 1 ml of culture medium containing 0.5% methylcellulose and incubated for 3 days at 37°C in 5% CO2 and plaques were developed and scored as described in the Method section. Mean percent inhibition +/- standard error of infection compared to the untreated control is plotted.

**Table 1:** IC_50_ values of Cysteamine HCL against wild type and major variants of SARS-CoV2. Inhibitory potency measured as IC_50_ was defined as the concentration of cysteamine-HCl that resulted in 50% reduction in the number of plaques compared to the untreated controls.

| Virus | IC<sub>50</sub> [mM] |
| --- | --- |
| wild type (Washington) | 1.252 |
| alpha variant | 1.528 |
| beta variant | 0.799 |
| gamma variant | 1.976 |
| lambda variant | 0.860 |
| delta variant (subvariant 1) | 1.066 |
| delta variant (subvariant 2) | 1.700 |
| delta variant (subvariant 3) | 1.373 |
| omicron variant (subvariant 1) | 0.671 |
| omicron variant (subvariant 2) | 1.006 |

**Table 1: IC<sub>50</sub> values of Cysteamine HCL against wild type and major variants of SARS-CoV2**
| **Virus** | **IC<sub>50</sub> [mM]** |
| --- | --- |
| wild type (Washington) | 1.252 |
| alpha variant | 1.528 |
| beta variant | 0.799 |
| gamma variant | 1.976 |
| lambda variant | 0.860 |
| delta variant (subvariant 1) | 1.066 |
| delta variant (subvariant 2) | 1.700 |
| delta variant (subvariant 3) | 1.373 |
| omicron variant (subvariant 1) | 0.671 |
| omicron variant (subvariant 2) | 1.006 |

To determine the duration of incubation of virus with cysteamine HCl required for maximal inhibition, the Delta variant was pre-incubated with either 5 or 10 mM concentration of cysteamine HCl for 0, 15, 30, 60 and 90 minutes. The mixture was then added to Vero-TMPRSS2 cells and infection continued for an additional 60 min resulting in a total association time of virus with cysteamine HCl of 60, 75, 90, 120 and 150 minutes. As shown in Figure 4, inhibition of delta variant was produced by both 5 and 10 mM cysteamine HCl under these conditions, with maximum inhibition at 10 mM when the virus and cysteamine HCl was incubated for 150 minutes. The level of inhibition of infection in this assay was slightly less than that observed in the dose response inhibition assay (Figure 3C), because the association time of virus with cysteamine HCl in this assay was 150 min when assayed compared to 180 min in Figure 3C.

**Figure 4.**
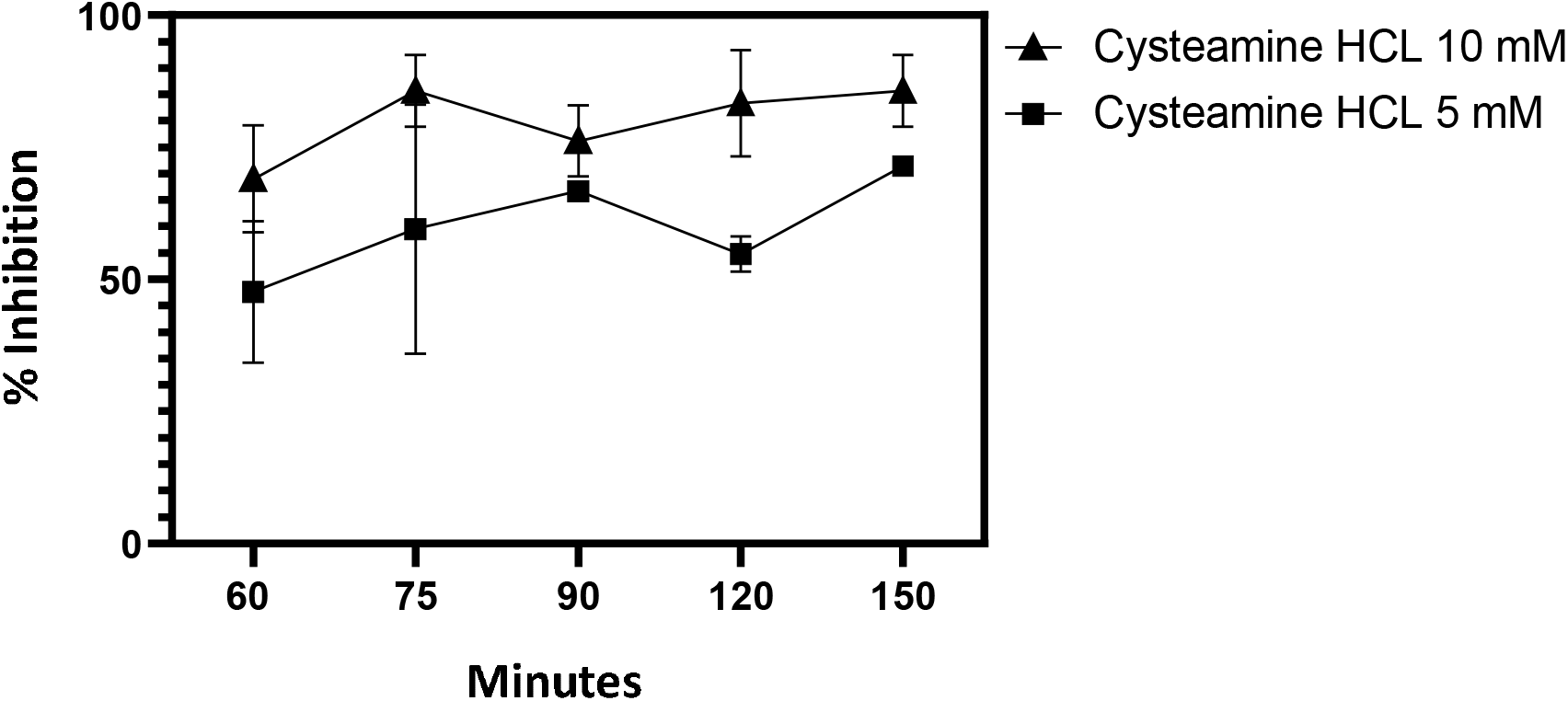
Kinetics of Inhibition of delta variant infection by cysteamine HCl. Delta variant (Subvariant 1) (37.5 pfu/ml) was preincubated with cysteamine HCl (5 or 10 mM) for 0, 15, 30, 60 and 90 min in a total volume of 600 μl of complete DMEM medium. The cysteamine HCl/virus mixture was then transferred to wells of Vero-TMPRSS2 cells in a total volume of 250 μl and incubated for 1 hour at 37°C in 5% CO2. Wells were then overlaid with 1 ml of culture medium containing 0.5% methylcellulose and plaques were developed as described in the Method section. In both assays plaques were developed and scored as described in the Method section. Mean percent inhibition of infection +/- standard error compared to the untreated control along with the total time of association of virus and cysteamine before addition of 0.5% methyl cellulose overlay was plotted.

## Discussion

In this study the antiviral activity of cysteamine HCl was assessed against wild type and major variants of SARS-CoV-2 that have emerged so far including the delta and omicron variants, which have spread widely throughout the world with devastating consequences. These assays were conducted using Vero-E6 cells which over-express the transmembrane serine protease II (Vero-TMPRSS2) because these cells are highly susceptible to infection with wild type and all variants of SARS-CoV2. Our results demonstrate *in vitro* antiviral activity of cysteamine against both wild type and multiple variants of SARS-CoV-2. Similar inhibition of the infectivity of SARS-CoV-2 with cysteamine was also demonstrated with IC_50_ values lower than what was noted in our study (14). This difference is most likely due to the highly susceptible target cells used in our study compared to the Vero E6 cells used in the previous work.

The spike protein of SARS-CoV-2 is reported to have fourteen disulfide bonds (19). It is likely that cysteamine reduces the disulfide bonds leading to altered conformation and instability of the receptor binding domain (RBD). This, in turn, inhibits binding of RBD with ACE-2 receptor on the target cells (19,20). Since disulfide residues in the RBD are conserved among the emergent variants, it is not surprising that all six variants were sensitive to inhibition by cysteamine HCl. Mutations affecting the disulfide bonds are predicted to denature the spike protein receptor binding site, leading to reduced virus infectivity and decreased reproduction rate. Therefore, any viable future mutations are predicted to remain susceptible to cysteamine (19).

Cysteamine is a free thiol with the odor and taste of rotten eggs, accounting for the olfactory and taste aversion to this medication. Despite this, cysteamine has been shown to be well tolerated when administered orally to patients with cystinosis. Side effects due to oral forms are primarily limited to nausea, vomiting and gastric hyperacidity. This limits compliance in some younger patients on chronic cysteamine therapy, however 94% of patients > 11 years of age reported always being compliant, compared to 50% < 11 years of age who were less (21). Cysteamine eye drops are well tolerated and are marketed in both immediate and long-acting forms. The first ocular form to be FDA approved, Cystaran, is recommended to be given hourly while awake. A more recent, long-acting form, Cystadrops, is administered four times a day while awake. Both are well tolerated but may provoke transient burning and mild eye irritation is some patients. Both are used in lifelong treatment to maintain the corneas free of cystine crystals.

It is accepted that nasal epithelium cells located in the nasopharynx are: (i) the initial source of individual infection with SARSCoV2; (ii) the location of rapid replication and mutation of the virus, and (iii) the primary source from which the virus spreads to others. The nasal epithelium is a target for SARSCoV-2 to enter and replicate via the concentrated ACE2 receptors on goblet cells (22). As the virus proliferates in the nasal epithelium, lysis of epithelial cells releases virions in a logarithmic progression which then transit to the trachea, bronchi and ultimately alveoli, leading to pneumonia, devastating illness (23), and markedly greater infectivity, as measured by R*_RI_* (24) Since cysteamine is currently marketed as two well tolerated ophthalmic preparations, Cystaran ^®^ (0.44%, = 56mM), and Cystadrops^®^ (0.37%, = 33mM) the drug could feasibly be administered topically to the nasal epithelium in the concentrations used for frequent ocular administration, and which are shown to inactivate the virus *in vitro* in tissue culture in highly permissive cells at concentrations between one tenth and one sixth that currently approved for chronic ophthalmic use by US FDA (see Figures 3 and 4). There it could act as a chemical impediment to nasal entry and thus to virus replication. Cysteamine employed as a nasal spray, cream or drops could function as both a preventative and mitigator of COVID-19 infection. The Delta variant is believed to result in significantly higher viral loads in the nasopharynx than the original virus, thus providing significantly greater opportunities for viral spread and mutation. Reducing or eliminating the virus’s ability to infect nasal epithelium cells by treatment with topical cysteamine would not only reduce its ability to sicken infected individuals, but also reduce or eliminate its ability to replicate, mutate and infect others. Nasal sprays to combat SARS Cov-2 have been proposed, including nasal administration of nitric oxide (NO) (25), carrageenan, Ivermectin, chloroquine, Niclosamide, steroids, ethyl lauroyl arginate, hypochlorous acid, povidone-iodine, antibodies, lipopeptide, and a PEGylated TLR2/6 agonist. All are in various levels of pre-clinical or clinical trials (26).

Effective utilization of a nasal spray or drops as a means of administering an antiviral agent to prevent or mitigate SARSCoV-2 infection requires coverage of the nasal epithelium. The nasal surface area is reported at 160 cm^2^ (27). Assuming a nasal spray of 0.5 ml in each nostril, a volume well tolerated by children (28), delivery of 1 ml total volume to the nasal surface would yield a potential uniform film thickness of the antiviral solution of 1 cm^3^ / 160 cm^2^ = 0.0063 cm, or 63 μ. This assumes both uniform distribution of the nasal spray throughout the nasal epithelium, and uniform distribution of ACE2 receptors, both of which are not known at this time. The dwell time of virus in the film may be increased by increasing the viscosity of the excipients, which may permit achieving dwell times up to a few hours. SARS-CoV-2 is inactivated in vitro in this study by cysteamine at 5 to 10 mM concentration, which is one sixth to one tenth the concentration approved by US FDA for chronic ophthalmic use in cystinosis. Nasal administration of pharmacologic agents is becoming increasingly favored, although most are used to deliver systemic agents or to target the CNS. As of this writing there are 3645 nasal administration results in PubMed.

## Summary

Results presented in this communication demonstrate significant inhibition of infection of variants of SARS-CoV-2 including delta, lambda and omicron variants by cysteamine HCl in a highly permissible cell line Vero-TMPRSS2. Cysteamine is a well-studied drug with very good safety profile with chronic use in patients with cystinosis. Administration of cysteamine as a topical agent to the nasal mucosa of both pre-exposed and infected individuals may be beneficial in number of ways via formation of a chemical barrier. It would thus prevent virus binding and inhibit infection of ACE-2 expressing epithelial cells, thereby reducing the amount of infectious virus entering the trachea-bronchial tree, and by reducing the amount of infectious virus produced in the nasal epithelium. This, in turn, may reduce the spread of virus through exhalation by the treated individuals.

## Acknowledgement

Cysteamine-HCL used in this study was provided by ACIC Pharmaceuticals Inc. (Ontario, Canada). We would like to thank Takaji Wakita from the National Institute of Infectious Diseases, Japan, for kindly providing the gamma variant and Mehul Suthar from the Emory University, USA for one of the subvariants of omicron used in this study. Rest of the virus stocks used here were obtained from BEI Resources, NIAID, NIH as mentioned in the Materials and Methods. We would like to thank Shelby O’Connor, John Baczenas and Corina Valencia from the University of Wisconsin-Madison, USA for deep sequencing our in-house expanded virus stocks and Adrian Creanga from the Vaccine Research Center-NIAID, USA, for the Vero TMPRSS2 cell line.

Jess Thoene and Robert Gavin are inventors on a patent application related to the use of cysteamine for the prevention and treatment of COVID 19 infections. It is assigned to the University of Michigan, licensed by ACIC Pharmaceuticals, Inc who funded this study.

## Notes

### Summary of Updates

This version now includes data showing inhibition of Lambda and Omicron variants.

## References

1. Prescott, L. F., Newton, R. W., Swainson, C. P., Wright, N., Forrest, A. R., and Matthew, H. (1974) Successful treatment of severe paracetamol overdosage with cysteamine. Lancet 1, 588–592

2. Thoene, J. G., Oshima, R. G., Crawhall, J. C., Olson, D. L., and Schneider, J. A. (1976) Cystinosis. Intracellular cystine depletion by aminothiols in vitro and in vivo. J Clin Invest 58, 180–189

3. Gahl, W. A., Reed, G. F., Thoene, J. G., Schulman, J. D., Rizzo, W. B., Jonas, A. J., Denman, D. W., Schlesselman, J. J., Corden, B. J., and Schneider, J. A. (1987) Cysteamine therapy for children with nephropathic cystinosis. N Engl J Med 316, 971–977

4. Smolin, L. A., Clark, K. F., Thoene, J. G., Gahl, W. A., and Schneider, J. A. (1988) A comparison of the effectiveness of cysteamine and phosphocysteamine in elevating plasma cysteamine concentration and decreasing leukocyte free cystine in nephropathic cystinosis. Pediatr Res 23, 616–620

5. Tenneze, L., Daurat, V., Tibi, A., Chaumet-Riffaud, P., and Funck-Brentano, C. (1999) A study of the relative bioavailability of cysteamine hydrochloride, cysteamine bitartrate and phosphocysteamine in healthy adult male volunteers. Br J Clin Pharmacol 47, 49–52

6. Gahl, W. A., Thoene, J. G., and Schneider, J. A. (2002) Cystinosis. N Engl J Med 347, 111–121

7. Gahl, W. A., Ingelfinger, J., Mohan, P., Bernardini, I., Hyman, P. E., and Tangerman, A. (1995) Intravenous cysteamine therapy for nephropathic cystinosis. Pediatr Res 38, 579–584

8. Bendel-Stenzel, M. R., Steinke, J., Dohil, R., and Kim, Y. (2008) Intravenous delivery of cysteamine for the treatment of cystinosis: association with hepatotoxicity. Pediatr Nephrol 23, 311–315

9. Mansouri, P., Farshi, S., Hashemi, Z., and Kasraee, B. (2015) Evaluation of the efficacy of cysteamine 5% cream in the treatment of epidermal melasma: a randomized double-blind placebo-controlled trial. Br J Dermatol 173, 209–217

10. Cherqui, S., and Courtoy, P. J. (2017) The renal Fanconi syndrome in cystinosis: pathogenic insights and therapeutic perspectives. Nat Rev Nephrol 13, 115–131

11. Barbouche, R., Miquelis, R., Jones, I. M., and Fenouillet, E. (2003) Protein-disulfide isomerase-mediated reduction of two disulfide bonds of HIV envelope glycoprotein 120 occurs post-CXCR4 binding and is required for fusion. J Biol Chem 278, 3131–3136

12. Anderson, R. A., Jr., Feathergill, K., Kirkpatrick, R., Zaneveld, L. J., Coleman, K. T., Spear, P. G., Cooper, M. D., Waller, D. P., and Thoene, J. G. (1998) Characterization of cysteamine as a potential contraceptive anti-HIV agent. J Androl 19, 37–49

13. Bergamini, A., Ventura, L., Mancino, G., Capozzi, M., Placido, R., Salanitro, A., Cappannoli, L., Faggioli, E., Stoler, A., and Rocchi, G. (1996) In vitro inhibition of the replication of human immunodeficiency virus type 1 by beta-mercaptoethylamine (cysteamine). J Infect Dis 174, 214–218

14. Khanna, K., Raymond, W., Jin, J., Charbit, A. R., Gitlin, I., Tang, M., Werts, A. D., Barrett, E. G., Cox,. M., Birch, S. M., Martinelli, R., Sperber, H. S., Franz, S., Pillai, S., Healy, A. M., Duff, T., Oscarson, S., Hoffmann, M., Pohlmann, S., Simmons, G., and Fahy, J. V. (2021) Thiol drugs decrease SARS-CoV-2 lung injury in vivo and disrupt SARS-CoV-2 spike complex binding to ACE2 in vitro. bioRxiv https://doi.org/10.1101/2020.12.08.415505

15. Alonzi, T., Aiello, A., Petrone, L., Najafi Fard, S., D’Eletto, M., Falasca, L., Nardacci, R., Rossin, F., Delogu, G., Castilletti, C., Capobianchi, M. R., Ippolito, G., Piacentini, M., and Goletti, D. (2021) Cysteamine with In Vitro Antiviral Activity and Immunomodulatory Effects Has the Potential to Be a Repurposing Drug Candidate for COVID-19 Therapy. Cells 2022, 11, 52. https://doi.org/10.3390/cells11010052

16. Chen, R. E., Zhang, X., Case, J. B., Winkler, E. S., Liu, Y., VanBlargan, L. A., Liu, J., Errico, J. M., Xie, X., Suryadevara, N., Gilchuk, P., Zost, S. J., Tahan, S., Droit, L., Turner, J. S., Kim, W., Schmitz, A. J., Thapa, M., Wang, D., Boon, A. C. M., Presti, R. M., O’Halloran, J. A., Kim, A. H. J., Deepak, P., Pinto, D., Fremont, D. H., Crowe, J. E., Jr., Corti, D., Virgin, H. W., Ellebedy, A. H., Shi, P. Y., and Diamond, M. S. (2021) Resistance of SARS-CoV-2 variants to neutralization by monoclonal and serum-derived polyclonal antibodies. Nat Med 27, 717–726

17. Edara VV, Manning KE, Ellis M, Lai L, Moore KM, Foster SL, Floyd K, Davis-Gardner ME, Mantus G, Nyhoff LE, Bechnak S, Alaaeddine G, Naji A, Samaha H, Lee M, Bristow L, Hussaini L, Ciric CR, Nguyen PV, Gagne M, Roberts-Torres J, Henry AR, Godbole S, Grakoui A, Sexton M, Piantadosi A, Waggoner JJ, Douek DC, Anderson EJ, Rouphael N, Wrammert J, Suthar MS. mRNA-1273 and BNT162b2 mRNA vaccines have reduced neutralizing activity against the SARS-CoV-2 Omicron variant. bioRxiv [Preprint]. 2021 Dec 22:2021.12.20.473557. doi: 10.1101/2021.12.20.473557. Update in: Cell Rep Med. 2022 Jan 24;3(2):100529. PMID: 34981056; PMCID: PMC8722593.

18. To, A., Wong, T. A. S., Lieberman, M. M., Thompson, K., Ball, A. H., Pessaint, L., Greenhouse, J., Daham, N., Cook, A., Narvaez, B., Flinchbaugh, Z., Van Ry, A., Yalley-Ogunro, J., Andersen Elyard, H., Lai, C. Y., Donini, O., and Lehrer, A. T. (2022) A Recombinant Subunit Vaccine Induces a Potent, Broadly Neutralizing, and Durable Antibody Response in Macaques against the SARS-CoV-2 P.1 (Gamma) Variant. ACS Infect Dis. 2022 Mar 9:acsinfecdis.1c00600. doi: 10.1021/acsinfecdis.1c00600. Epub ahead of print. PMID: 35263081; PMCID: PMC8938837.

19. Grishin AM, Dolgova NV, Landreth S, Fisette O, Pickering IJ, George GN, Falzarano D, Cygler M. Disulfide Bonds Play a Critical Role in the Structure and Function of the Receptor-binding Domain of the SARS-CoV-2 Spike Antigen. J Mol Biol. 2022 Jan 30;434(2):167357. doi: 10.1016/j.jmb.2021.167357. Epub 2021 Nov 12. PMID: 34780781; PMCID: PMC8588607.

20. Yan, R., Zhang, Y., Li, Y., Xia, L., Guo, Y., and Zhou, Q. (2020) Structural basis for the recognition of SARS-CoV-2 by full-length human ACE2. Science 367, 1444–1448

21. Ariceta, G., Lara, E., Camacho, J. A., Oppenheimer, F., Vara, J., Santos, F., Munoz, M. A., Cantarell, C., Gil Calvo, M., Romero, R., Valenciano, B., Garcia-Nieto, V., Sanahuja, M. J., Crespo, J., Justa, M. L., Urisarri, A., Bedoya, R., Bueno, A., Daza, A., Bravo, J., Llamas, F., and Jimenez Del Cerro, L. A. (2015) Cysteamine (Cystagon(R)) adherence in patients with cystinosis in Spain: successful in children and a challenge in adolescents and adults. Nephrol Dial Transplant 30, 475–480

22. Sungnak, W., Huang, N., Becavin, C., Berg, M., Queen, R., Litvinukova, M., Talavera-Lopez, C., Maatz, H., Reichart, D., Sampaziotis, F., Worlock, K. B., Yoshida, M., Barnes, J. L., and Network, H. C. A. L. B. (2020) SARS-CoV-2 entry factors are highly expressed in nasal epithelial cells together with innate immune genes. Nat Med 26, 681–687

23. Hou YJ, Okuda K, Edwards CE, Martinez DR, Asakura T, Dinnon KH 3rd, Kato T, Lee RE, Yount BL, Mascenik TM, Chen G, Olivier KN, Ghio A, Tse LV, Leist SR, Gralinski LE, Schäfer A, Dang H, Gilmore R, Nakano S, Sun L, Fulcher ML, Livraghi-Butrico A, Nicely NI, Cameron M, Cameron C, Kelvin DJ, de Silva A, Margolis DM, Markmann A, Bartelt L, Zumwalt R, Martinez FJ, Salvatore SP, Borczuk A, Tata PR, Sontake V, Kimple A, Jaspers I, O’Neal WK, Randell SH, Boucher RC, Baric RS. SARS-CoV-2 Reverse Genetics Reveals a Variable Infection Gradient in the Respiratory Tract. Cell. 2020 Jul 23;182(2):429–446.e14. doi: 10.1016/j.cell.2020.05.042. Epub 2020 May 27. PMID: 32526206; PMCID: PMC7250779.

24. Ito K, Piantham C, Nishiura H. Predicted dominance of variant Delta of SARS-CoV-2 before Tokyo Olympic Games, Japan, July 2021. Euro Surveill. 2021 Jul;26(27):2100570. doi: 10.2807/1560-7917.ES.2021.26.27.2100570. PMID: 34240695; PMCID: PMC8268651.

25. Winchester, S., John, S., Jabbar, K., and John, I. (2021) Clinical efficacy of nitric oxide nasal spray (NONS) for the treatment of mild COVID-19 infection. J Infect 83, 237–279

26. Pilicheva B, Boyuklieva R. Can the Nasal Cavity Help Tackle COVID-19? Pharmaceutics. 2021 Oct 3;13(10):1612. doi: 10.3390/pharmaceutics13101612. PMID: 34683904; PMCID: PMC8537957.

27. Illum, L. (2000) Transport of drugs from the nasal cavity to the central nervous system. Eur J Pharm Sci 11, 1–18

28. Tsze, D. S., Ieni, M., Fenster, D. B., Babineau, J., Kriger, J., Levin, B., and Dayan, P. S. (2017) Optimal Volume of Administration of Intranasal Midazolam in Children: A Randomized Clinical Trial. Ann Emerg Med 69, 600–609

